# Molecular architecture of the *Legionella* Dot/Icm type IV secretion system

**DOI:** 10.1101/312009

**Authors:** Debnath Ghosal, Yi-Wei Chang, Kwang Cheol Jeong, Joseph P. Vogel, Grant J. Jensen

## Abstract

*Legionella pneumophila* survives and replicates inside host cells by secreting ~300 effectors through the Dot/Icm type IVB secretion system (T4BSS). Understanding this machine’s structure is challenging because of its large number of components (27) and integration into all layers of the cell envelope. Previously we overcame this obstacle by imaging the Dot/Icm T4BSS in its native state within intact cells through electron cryotomography. Here we extend our observations by imaging a stabilized mutant that yielded a higher resolution map. We describe for the first time the presence of a well-ordered central channel that opens up into a windowed large (~32 nm wide) secretion chamber with an unusual 13-fold symmetry. We then dissect the complex by matching proteins to densities for many components, including all those with periplasmic domains. The placement of known and predicted structures of individual proteins into the map reveals the architecture of the T4BSS and provides a roadmap for further investigation of this amazing specialized secretion system.

## Introduction

Bacterial cells have evolved a variety of specialized nanomachines to secrete proteins, nucleic acids, and other materials into their environment^1,2^. Among these, the type IV secretion system (T4SS) is arguably the most complex and versatile. T4SSs include the well-studied conjugation machine that *Escherichia coli* and other bacteria use to transfer plasmids from donor to recipient cells, as well as the *Cag* T4SS from *Helicobacter pylori* that injects a carcinogenic effector protein into human intestinal epithelial cells^3,4^. Based on component number and similarity, T4SSs are classified into three types: IVA, IVB, or other^5,6^. The archetype type IVA system is the VirB/D4 T4ASS, widely used to genetically engineer plants, that *Agrobacterium tumefaciens* uses to inject bacterial genes into plant roots, generating tumors^7,8^. It consists of a lytic transglycosylase (VirB1), pilins (VirB2 and VirB5), inner membrane (IM) proteins (VirB3, VirB6, VirB8), ATPases (VirB4, VirB11 and VirD4) and a three-component subcomplex (VirB7, VirB9, VirB10) that spans the inner and outer membranes^9^.

Type IVB systems, represented by the IncI conjugative plasmids (R64 and ColIbP9) and the Dot/Icm system, are more complex than T4ASSs^9,10^. The Dot/Icm (<u>D</u>efective in <u>o</u>rganelle <u>t</u>rafficking/<u>I</u>ntracellular <u>m</u>ultiplication) T4BSS enables *Legionella pneumophila*, the causative agent of Legionnaires’ disease, to secrete ~300 different effector proteins into human and other host cells, allowing the pathogen to survive and replicate within phagocytic compartments^11,12^. It has ~27 components, including an outer membrane (OM) protein (DotH), OM lipoproteins (DotC, DotD, DotK), a periplasmic protein (IcmX), IM proteins (DotA, DotE, DotF, DotG, DotI, DotJ, DotM, DotP, DotU, DotV, IcmF, IcmT, and IcmV), IM-associated ATPases (DotB, DotL, DotO), and soluble cytosolic proteins (DotN, IcmQ, IcmR, IcmS, IcmW, and LvgA)^13^. The ATPases of the Dot/Icm and VirB/D4 systems belong to the same classes^13^, but the only other clear sequence similarity between T4ASS and T4BSS components occurs within the C-terminus of DotG, which matches part of the VirB10 sequence^13,14^

T4SSs are dynamic multi-protein machines that, in Gram-negative bacteria, span and interact with two cellular compartments (cytoplasm and periplasm), two membranes (inner and outer) and a peptidoglycan cell wall, making purification and structural analysis challenging. Important progress has been made by solving the structures of some components and subcomplexes^15–20^, but it is not always clear how the individual pieces fit into the whole or into the context of the cell envelope. Previously, we showed that these difficulties can be overcome by imaging T4SSs in their native state within intact cells by electron cryotomography (ECT)^14,21^. By identifying, aligning, and averaging hundreds of individual Dot/Icm particles, we produced a sub-tomogram average revealing the overall structure of the machine, including several distinct densities^14^. Here we identify a tagged component that stabilizes the structure, improving the sub-tomogram average and revealing higher-resolution details of the complex. We then dissect the structure by imaging mutants lacking components or expressing tagged versions, locating all 10 OM and periplasmic components. Based on known structures of components and new predictions, we then build an “architectural” map of the Dot/Icm T4BSS with implications for the function, assembly, and evolution of this complex machine.

## Results

### Structural details of the Dot/Icm T4BSS

Previously, we identified Dot/Icm T4BSSs in electron cryotomograms of wild-type *L. pneumophila*. Since individual tomograms are noisy (due both to dose limitations and to the cellular context, with random, freely-diffusing proteins in the vicinity of the T4BSS), we generated a sub-tomogram average to reveal only those densities consistently present in the machine. Since the T4BSS is flexible, we calculated and combined independent averages, aligning the particles first on the OM-associated densities and then on the cytoplasmic densities (Fig. 1A)^14^.

**Figure 1.**
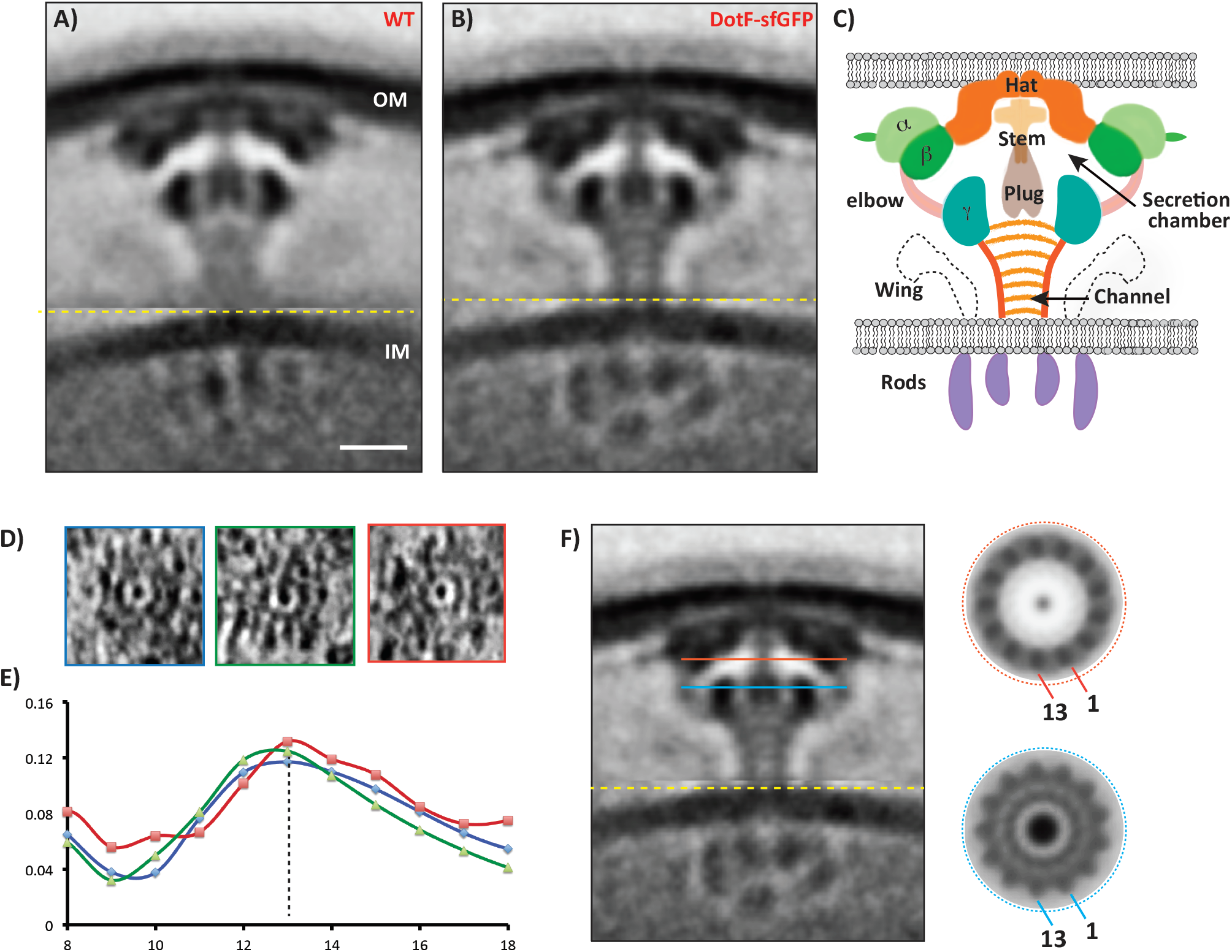
Overall structure of the *dot/icm* T4BSS. Central tomographic slices through the (A) wild-type and (B) DotF-sfGFP sub-tomogram averages, showing the improved resolution of the DotF-sfGFP structure. Note that because Dot/Icm particles are flexible, all the sub-tomograms going into the average were first aligned on the OM-associated densities and then on the IM-associated densities, separately; the image shown is a composite of the two averages concatenated at the yellow line. Scale bar 10 nm. C) Schematic of the major densities in the structure, named for reference. D) Tomographic slices through individual particles showing top views. E) Rotational cross-correlation coefficients of the three top-view particles for symmetries from 8-to 18-fold, showing that 13-fold was the strongest in each case. F) Applying 13-fold rotational symmetry at the levels of the red and blue lines in the DotF-sfGFP average produced clear structures (other symmetries failed to produce regular density patterns). For scale, the diameters of the red and blue circles are both 32 nm.

We repeated this process with a strain expressing DotF tagged with superfolder GFP (DotF-sfGFP), which we found remarkably stabilized the particles, resulting in a higher-resolution average of the complex (Fig. 1B, Fig. S1). All the previously identified features, including the alpha, beta, gamma, hat, stalk, wing, rod and stem densities, as well as the globular density in the middle of the gamma ring that we now term the plug, were also visible in the DotF-sfGFP structure. In addition, we saw that the stalk is not a solid object, but rather an ~14 nm-long funnel-shaped channel with thick (~2 nm) walls and a central lumen ~4 nm wide at the base (adjacent to the IM) (Fig. 1B, C). The lumen of the channel was not empty; we saw diffuse density within and a few thin striations perpendicular to the channel axis (Fig. 1B, C).

In the DotF-sfGFP reconstruction, there was a clear connection between the beta and the gamma densities (labelled “elbow”), revealing that the stalk channel leads into a large (~32 nm wide) secretion chamber surrounded by the hat, beta, elbow, and gamma rings. The design is therefore reminiscent of the T4ASS core-complex where the O-layer and the I-layer together form a secretion chamber^19^. Noting that the elbow density was much less pronounced than the rings, it is likely not continuous around the circumference of the chamber, but is rather a series of arm-like densities separated by gaps allowing access to the periplasm. Finally, the wings were also more pronounced in the DotF-sfGFP map, and there was a thin additional ring of density, hinted at in the wild-type map but not as clear, protruding out of the alpha density parallel to the OM (Fig. 1B, C).

In top views of individual particles, we could count 13 subunits around the ring (Fig 1D). To confirm this symmetry, we calculated rotational cross-correlations of slices through several individual particles at the level of the alpha and beta densities. The peak was consistently 13-fold (Fig. 1E). In the average, clear features emerged at the level of the alpha/beta and gamma densities when 13-fold, but not other, symmetry was applied (Fig. 1F). We conclude that the alpha, beta, and gamma rings are 13-fold symmetric.

The cytoplasmic densities were also better resolved in the DotF-sfGFP strain. A central slice through the average exhibited two shorter central rods with globular densities below each flanked by two longer rods (Fig. S2), but slices in front of or behind that central slice (displaced in a direction perpendicular to the axis of the complex) exhibited four longer parallel rod-like densities (Fig. S2). These patterns were reminiscent of the cytoplasmic complex of the *H. pylori* Cag T4SS, though not exactly the same, and suggest the presence of a short central barrel on the machine’s axis surrounded by taller barrels around the periphery^21^. An additional density resides directly below the short central barrel (Fig. 1B, Fig. S2A).

### Component dissection of the Dot/Icm T4BSS

Knowing the overall structure of the T4BSS, we next sought to locate components within the structure. To do that, we imaged a series of *L. pneumophila* mutants with individual or combinations of components deleted, or with components tagged with sfGFP to provide additional density (Fig. 2). We confirmed the absence of protein expression in the various deletion mutants and the presence of the sfGFP fusions by Western blot analysis (Fig. S3). Some mutants failed to produce visible T4BSSs; for those that did (insets in Fig. 2), we calculated sub-tomogram averages. The number of tomograms collected for each of the mutants and the number of particles used for sub-tomogram averaging are listed in SI Table 1.

**Figure 2.**
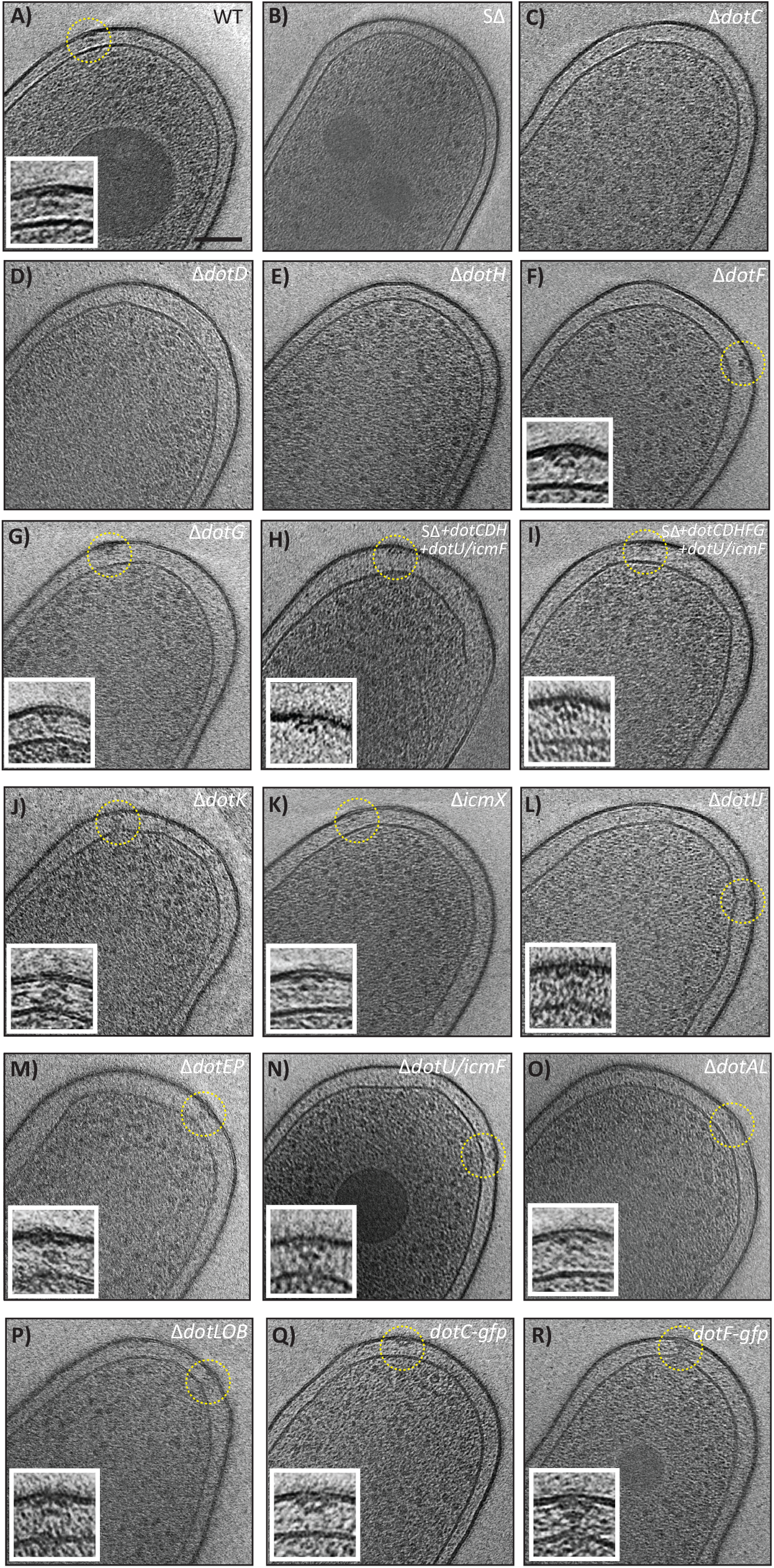
Electron cryotomography of intact *Legionella pneumophila* cells expressing T4SSs. Panels show tomographic slices 8-nm thick through one representative cell of each strain imaged. The strain identity is given in the upper right corner of each panel. In those strains in which T4SS particles or sub-particles were seen, an enlarged image of an example particle is shown in the inset. Scale bar 100 nm.

As expected, no T4BSS particles were visible in a “super-deletion” (SΔ) strain in which all of the *dot/icm* genes are deleted (Fig. 2B)^14^. Next we imaged strains missing each of the proteins of the previously described core-transmembrane complex^22^. No particles were found in the *ΔdotC, ΔdotD*, or *ΔdotH* strains, revealing that all three are critical for the initiation of assembly, but particles were seen in the *ΔdotF* and *ΔdotG* strains (Fig. 2C-G). Based on two independent analysis of the Dot/Icm apparatus^15,22,23^, we analyzed SΔ strains expressing the core-transmembrane subcomplex (DotDCHGF) versus a smaller complex of only DotDCH in the presence of the polar targeting factors DotU and IcmF. In both cases, we observed particles that resembled portions of the T4BSS structure (Fig. 2H, I). Finally, we imaged nine other strains with individual or sets of components deleted (Fig. 2J-P) or components tagged with sfGFP (Fig. 2Q,R).

For each mutant strain that exhibited T4BSS particles, we calculated a difference map between its sub-tomogram average and that of the wild-type (Fig. 3). In each case clear differences were apparent. In some cases, discussed below, loss of a component blurred the average, reflecting destabilization of the complex.

**Figure 3.**
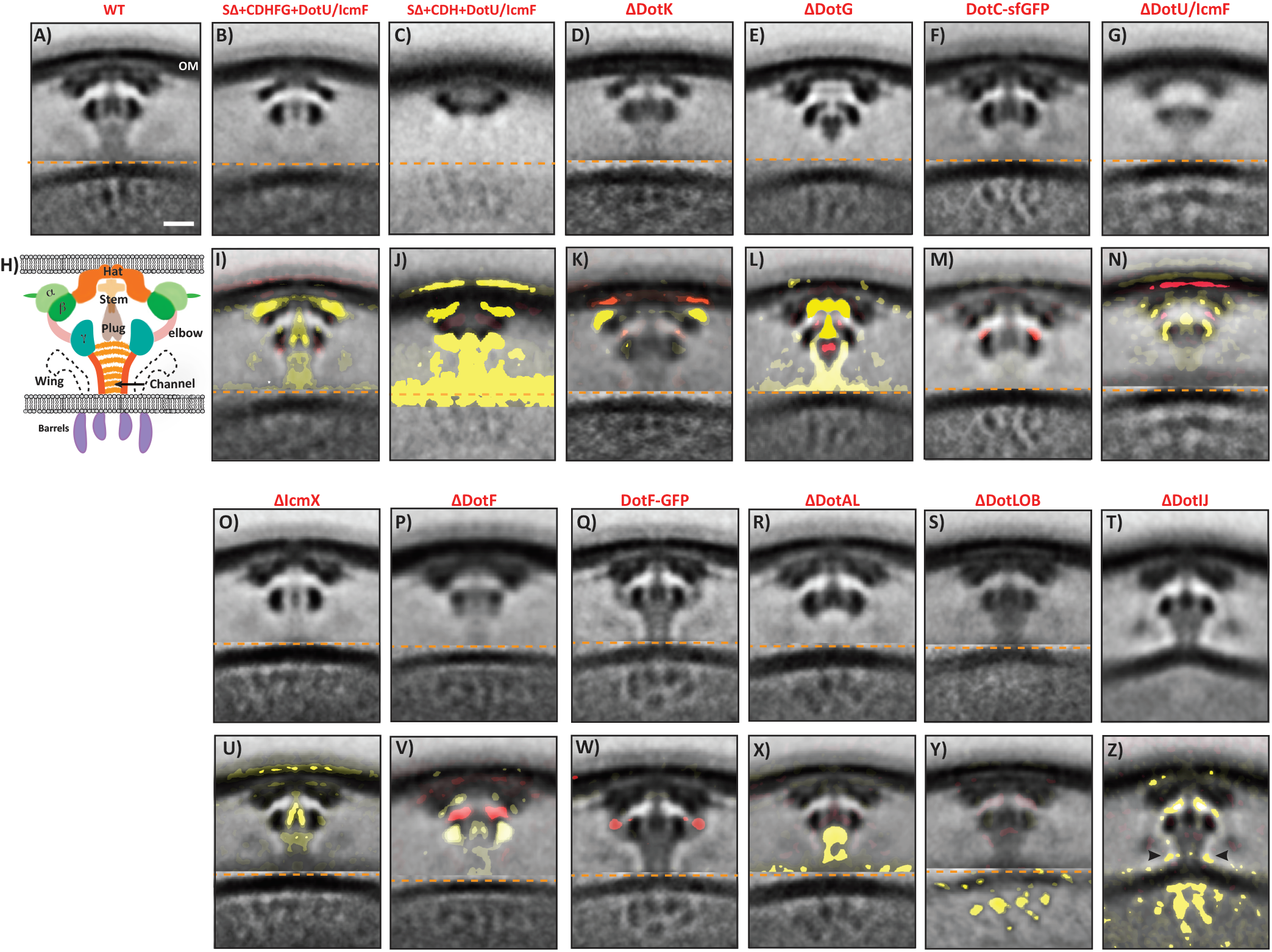
Mutant structures and difference maps. Top rows: Central slices through the sub-tomogram average structures of each strain imaged. Bottom rows: Central slices through the difference maps comparing each average to the wild-type. Yellow represents missing densities and red extra densities. Weak to strong intensities correspond to density differences from one to three standard deviations, respectively, overlaid on the mutant sub-tomogram average. Note isolated differences (as in panels M, U, and W) are candidate locations of missing or additional densities; matched yellow/red pairs on either side of a structure (as in panel V) likely indicate movements. Note in the case of *dotIJ*, the full image is from the IM-aligned averages since the feature of interest is at the same level as the concatenation. Scale bar 10 nm.

### Assigning locations of Dot/Icm T4BSS components

The sub-tomogram averages and difference maps of the mutants allowed us to place components into the overall structure (Fig. 4), guided by existing structural knowledge and predictions of secondary and tertiary structure (Fig. S4, Fig. S5). Here we summarize the process for each component.

**Figure 4.**
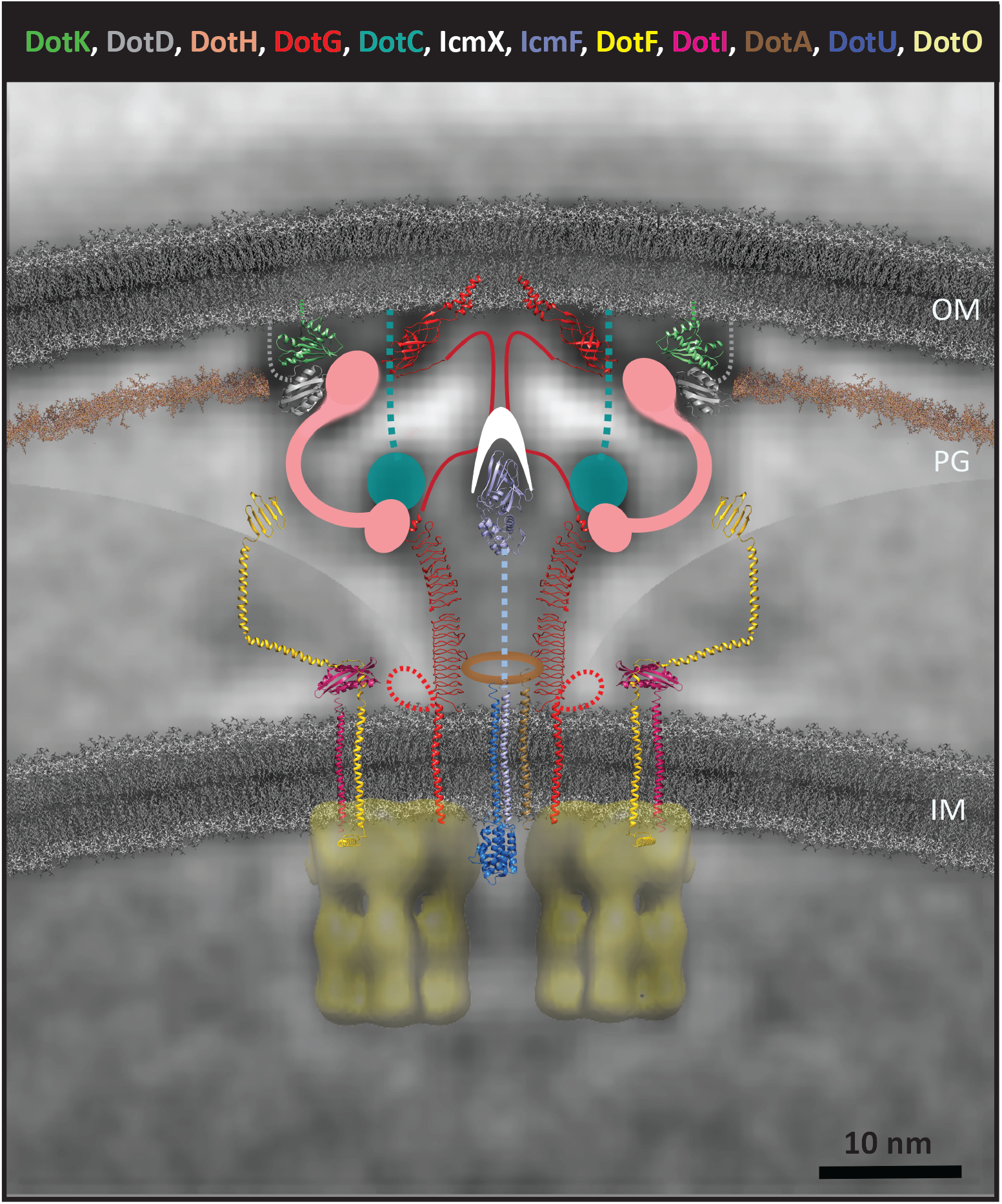
Architectural model of the Dot/Icm T4BSS. Based on the difference maps and evidence discussed in the text, known and predicted structures of T4BSS components are superimposed on the central slice of the DotF-sfGFP average. The wing region is from an average of particles aligned on that area, so that parts of three different averages are shown. Note that the relative orientation of the component structures are not known – the purpose of this schematic is simply to show where in the T4BSS each component is located and how its size and shape compare to the ECT densities. Components whose structures are not known or confidently predictable are depicted as circles (DotC) or as the shape of densities seen in the sub-tomogram averages or difference maps (DotH and IcmX). The low-resolution cryo-EM single particle reconstruction of VirB4 is used to represent two peripheral cytoplasmic ATPases (DotO). Presumably-extended polypeptide links to the OM are shown as dotted lines. Sequences in DotG with unknown structure are shown as solid lines with speculative paths drawn only to remind viewers that they may have important functions. Many other transmembrane helices are present that are not shown – so many in fact that they likely pack this region of the IM: DotA for instance is predicted to have 8-11 transmembrane helices and exist within the T4BSS as a dodecamer, and five additional proteins predicted to be almost entirely encompassed within the IM (DotE, DotP, DotV, IcmT, and IcmV) are not shown. Note other cytoplasmic proteins in the system are also not shown since we do not know yet where to place them. OM = outer membrane, PG = peptidoglycan cell wall, IM = inner membrane. Lipids are shown in grey and peptidoglycan in brown. Scale bar 10 nm.

#### DotK

DotK is highly conserved among *Legionella* and *Coxiella* species^24^, but deletion of the gene results in only a partial growth defect in protozoan hosts, and no defect for growth within macrophages^11,25^. In the *ΔdotK* average, all densities were present except alpha (Fig 3D, K). DotK is part of the OmpA domain family (Pfam F00691)^24^, bacterial peptidoglycan binding proteins that include the OM porin OmpA, the flagellar protein MotB, and peptidoglycan-associated proteins (PALs). Residues 2-131 are predicted to adopt the same fold as the C-terminal peptidoglycan binding domain of *Pseudomonas aeruginosa* OprF (PDB: 5U1H) from *Pseudomonas aeruginosa* (Fig. S4; Phyre2 server 100% confidence, 21% identity). This structure fit the alpha density well (Fig. 4), immediately adjacent to the thin line of density parallel to the OM, which we interpret to be ordered peptidoglycan.

#### DotG

DotG is the only protein in the T4SS superfamily with a clear sequence homolog in every subfamily (VirB10/TrbI in the T4ASS, TraO in the IncI plasmid-like systems, and DotG/IcmE in the T4BSS). The conserved part of the protein is the C-terminal TrbI domain (Pfam PF03743, residues 844–1045) (Fig. S4). The structure of this domain was solved in complex with the C-termini of VirB7 and VirB9^20^. The TrbI domain was seen to form a 14-fold-symmetric dome-shaped bowl with a central hole lined by two alpha helices per monomer thought to penetrate the OM^20^. We previously speculated that the TrbI domain of DotG forms a similar dome just underneath the OM in the T4BSS system as well, and showed that the size and shape of the TrbI domain matched the hat density in the Dot/Icm structure^14^. In the *ΔdotG* average, the hat was missing (Fig. 3E, L). Placing the TrbI domain structure into the hat density of the Dot/Icm complex in the orientation proposed in current models^19,20^ fit well (Fig. 4). The *ΔdotG* structure showed additional differences: the stem and stalk were also missing, the gamma ring was distorted, and the plug was several nanometers closer to the IM (Fig. 3E). To understand these differences, we considered the N-terminal sequence of DotG. Interestingly, T4BSS DotG-family proteins are significantly longer than the VirB10-family proteins due to the presence of an ~600 residue variable region containing repeats^11^. We found that both the *Legionella* and *Coxiella* DotG repeat regions are predicted to fold into long β-helices like those found in various bacterial effector proteins (Phyre2 99.9% confidence matches to PDB 3NB2 and 3DU1) (Fig. S4). The length, width, and even slight curvature of the predicted β-helices matched the apparent length, width, and curvature of the stalk channel wall, and 13 copies of the predicted β-helix structure could form a ring with the same inner and outer dimensions as the channel (Fig. S6C).

Because the stem was missing and the plug displaced in the *ΔdotG* structure, we speculate that the ~100 residues between the TrbI and β-helix domains form the stem and interact with the plug enroute to the stalk channel. Finally, it is unclear if the ~160 residues between the repeats and the N-terminal transmembrane helix are ordered or large enough to be seen in the averages, but they may form part of and/or interact with the wings (see below).

#### DotC

Recent immunofluorescence imaging shows that DotC interacts with DotH^23^, and earlier cellular fractionation and coimmunoprecipitation studies revealed that DotC/DotD/DotH form a subcomplex that associates DotH with the outer membrane^15,22^, but little else is known about DotC. No particles were seen in the *ΔdotC* mutant (Fig. 2C), but the DotC-sfGFP sub-tomogram average and difference map revealed additional density at the top of the gamma density (Fig. 3F, M). Consistent with this, the gamma density was one of only two densities (with beta) seen in the SΔ + DotCDH + DotU/IcmF map (Fig. 3C, Fig. S7). This subcomplex was heterogeneous, but it was clear that the beta and gamma and part of the plug densities were present (Fig. 3C and J). In the average, the gamma density appeared smaller than in wild-type, but gamma densities appeared full-sized in individual particles (Fig. S7), revealing that it is present, but more mobile. Unfortunately, the structure of DotC could not be predicted, so we estimated its size: a minimum of 29 residues would be required to span the ~10 nm distance from the membrane to the top of gamma. The 4.4 nm diameter of the rest of the protein (assuming a spherical assembly of the remaining 255 residues and 1.3g/mL average protein density) is less than the length of the gamma density, so we represented it as a circle at the top of gamma (Fig. 4).

#### DotH

DotH is associated with the OM, though it is not a lipoprotein^22^. Immunofluorescence and cell fractionation studies show that DotH binds DotC, and this complex is then bound by DotD, which causes DotH to become associated with the outer membrane^15,22,23^. Consistent with the immunofluorescence data^23^, no T4BSS particles were observed in the *ΔdotH* mutant. Unfortunately our attempts to generate a functional DotH-sfGFP were unsuccessful. The subcomplex of DotCDH + DotU/IcmF, however, exhibited the beta and gamma rings and the plug (Fig. 3C). Having located DotC in the top of the gamma ring, and with evidence that IcmF is in the plug (see below), we assumed the beta ring and bottom of the gamma ring were occupied by DotH and DotD. (Note DotU cannot be the beta or gamma ring, since it has only a few periplasmic residues.)

While not a clear match, the highest-scoring prediction for DotH structure from Phyre2 was VirB9, based on similarities between residues 168-227 of DotH and a model of the N-terminal domain of VirB9 (49% confidence). The C-terminal domain of VirB9 is known to bind the periphery of the TrbI domain of VirB10 in the O-layer and the N-terminus forms another domain in the lower I-layer^20,26^. Prediction of DotH structure using various programs (I-TASSER, Phyre2 and Quark) also suggested that DotH included two or more separate domains (Fig. S8). We therefore propose that DotH forms i) the central part of beta (where a domain roughly the size of the C-terminus of VirB9 might bind DotG), ii) the bottom of gamma (where another domain might bind DotC), and iii) the elbow between them (Fig. 4, Fig. S8C).

#### DotD

DotD is an OM lipoprotein attached to the membrane at cysteine 19. Following a relatively unstructured 47-residue linker, DotD contains a folded C-terminal N0 secretin domain whose structure has been solved^15^. Because another N0 domain was recently found in *Xanthomonas citri* VirB7, and it too is preceded by a short linker and an OM-attached cysteine, it was proposed that DotD is a VirB7 homolog^27^ A crystal structure of the *E. coli* pKM101 T4SS outer membrane complex consisting of parts of VirB7, VirB9 and VirB10 showed that VirB7 lines the tops of VirB9 and VirB10 immediately adjacent to the OM, but the *E. coli* VirB7 lacks a C-terminal N0 domain^20^. In order to investigate the location of the N0 domain in the *X. citri* VirB7, Souza et al. solved an NMR structure of the interface between the *X. citri* VirB9 and the VirB7 linker, and then used molecular dynamics to dock the N0 domain, which placed the N0 domain around the periphery of the VirB9 O-layer (Fig S9A, C)^27^. We therefore assumed that just as in *E. coli* and *X. citri* VirB7s, 26 residues of the DotD linker line the top of DotG (the hat) and DotH (the central part of beta). The linker in DotD contains ~10 additional residues so we placed it in the only unaccounted-for density within 3.5 nm (corresponding to an extended polypeptide chain of 10 residues): the periphery of beta, just where the *X. citri* N0 domain has been predicted to bind (Fig. S9, Fig. 4). Because there were no T4BSS particles in the ΔdotD mutant and our attempts to generate a functional DotD-sfGFP were unsuccessful, we could not directly confirm our mapping of DotD.

#### DotU/IcmF

DotU and IcmF are both integral IM proteins partially required for the intracellular growth of *L. pneumophila*^28^. Phylogenetic and sequences analyses have established that DotU and IcmF are homologs of the type VI secretion system TssL and TssM^28,29^. In a *ΔdotU ΔicmF* double mutant, the abundances of DotH, DotF and DotG are all reduced in late stationary phase, which led to the proposal that DotU and IcmF play an essential role in stabilizing the Dot/Icm core-complex^28^. Jeong et al. recently found that DotU and IcmF are the polar localization factors that recruit DotCH complexes to the poles to initiate assembly^23^.

In the *ΔdotU ΔicmF* double deletion mutant (Fig. 2N), we found polar as well as nonpolar particles, confirming that loss of DotU/IcmF results in mislocalization of the T4BSS. The average of these complexes exhibited low resolution (Fig. 3G), indicating that in addition to recruiting the T4BSS, DotU/IcmF also stabilizes it, suggesting a structural role. The difference map revealed a decrease in the plug density and a decreased diameter of the gamma ring (Fig. 3G, N). The structures of both the cytoplasmic domain of DotU and a periplasmic domain of IcmF (residues 738-972) are known by homology to TssL and TssM^30,31^. Only the C-terminus of IcmF could reach far enough into the periplasm to contact DotCH in the gamma and beta rings. The periplasmic domain of IcmF fits the plug density, so we propose the C-terminus of IcmF may form part of the plug (Fig. 4). This is consistent with the reduced diameter of the gamma ring in the absence of IcmF and would explain why it stabilizes the rest of the T4BSS, and how it recruits DotCH.

#### IcmX

IcmX is a soluble periplasmic protein^13,22^ conserved throughout *Legionella, Coxiella* and *Rickettsiella*, and a Δ*icmX* mutant exhibits a severe intracellular growth defect^32^. In *Legionella* and *Coxiella*, a truncated form of IcmX (residues ~165-466) is released in the culture media in a T4BSS-dependent manner^33,34^, but translocation across eukaryotic cell membranes has not been detected^33^. In the *E. coli* F-plasmid system, the counterpart of IcmX, TraW, is essential for F-pilus assembly^35^. Here we found that in the absence of IcmX, the T4BSS still assembled (Fig. 2K), but the periphery of the plug was missing (Fig. 3O, U), leading us to propose that IcmX is in this location. IcmX’s position in the channel rationalizes how it can be released.

#### DotF

DotF has been suggested to interact with, and regulate the activity of, DotG in the IM^22^. Several Dot/Icm substrates have been reported to interact with DotF, although this interaction has been called into question^36,37^, and *ΔdotF* mutants show partial growth defects in protozoan cells and macrophages^36^. Comparing the two reconstituted subcomplexes (SΔ + DotCDHFG + DotU/IcmF) and (SΔ + DotCDH + DotU/IcmF), we found that the former, with two added components (DotF and DotG), had two additional densities: the hat region we have now identified as the C-terminus of DotG, and the wings. This suggests the wings are DotF. Confirming the location of DotF, we saw that the wings were missing in the *ΔdotF* mutant (Fig. 3P,V, note that loss of the light wing densities was not highlighted by the difference map). Deletion of DotF did not affect assembly of the rest of the complex (Fig. 2F), but it did affect its stability, as evident from the low resolution sub-tomogram average (Fig. 3P). The difference map showed losses of density below and gains of density above the gamma ring, indicating that the gamma ring drifts toward the OM in the absence of DotF (Fig. 3V).

In many individual wild-type particles, wing densities were just as visible as alpha, beta, or gamma densities, but their positions varied between particles, washing them out in the average (Fig. S10A-I). In addition, the alpha, beta and gamma densities formed complete rings, making them appear even darker relative to the wings. We therefore computed additional sub-tomogram averages aligning particles with a mask centered on the wing region (from the IM up to and including the bottom of the gamma ring), which enhanced the density of the wings. In the *ΔdotG* structure the wing density was weaker but still visible, and it was completely absent in the *ΔdotF* structure, supporting the notion that the wings are largely DotF, stabilized by DotG (Fig. S10J-N). To further test this, we imaged a C-terminal sfGFP fusion of DotF (Fig. 3Q, W). In addition to stabilizing the entire complex (see above), there was a new density at the joint of the elbow between the gamma and beta densities, just above the wings, implying that the C-terminal domain of DotF is present at the joint of the elbow.

Phyre2, I-TASSER, and Quark prediction suggested that DotF folds into a long, potentially jointed alpha-helical structure capped by a C-terminal domain similar to that of PilP/GspC. We therefore represented DotF as a flexible arm above the peripheral barrels that can reach to the elbow of beta/gamma (see Discussion) (Fig. 4).

#### DotA

DotA is released into the culture media in a Dot/Icm-dependent manner, where it forms ring-like structures of uniform 10-nm diameter^33^. Its IncI plasmid homolog, TraY, has been described as an “extended-VirB6” protein located at the substrate transfer channel, suggesting a broadly conserved role^38^. In *L. pneumophila*, the *ΔdotA* mutant results in a nonfunctional Dot/Icm machine and exhibits an intracellular growth defect^33,39,40^.

To locate DotA, we examined the *ΔdotA ΔdotL* average. We reasoned that since DotL is a cytoplasmic ATPase (and in the *ΔdotL ΔdotO ΔdotB* deletion strain, the OM and periplasmic densities remained essentially unchanged (see below, Fig. 3S, Y)), any structural change in the OM and/or periplasmic complexes in the *ΔdotA ΔdotL* mutant would be because of the absence of DotA. The biggest difference was a major density missing in the top of the stalk channel (Fig. 3R, X), with diminished density in the stalk channel walls in that region. Since the diameter of the stalk roughly matched the diameter of purified DotA rings, we therefore propose that the periplasmic domain of DotA^41^ is located in the top of the stalk channel, where it stabilizes the walls and can be released (Fig. 4).

#### DotL/DotO/DotB

DotO is an IM-associated ATPase containing Walker A and Walker B motifs and distantly related to the VirB4 ATPase of the T4ASS^13^. DotO/VirB4 ATPases are the most conserved component of the T4SS superfamily^42^. VirB4 has been shown to form IM-associated hexameric barrels beneath the T4ASS complex^19^. DotL, a member of the type IV coupling protein family (T4CP) and a distant homolog of the VirD4 ATPase of the T4ASS^16^, is structurally related to the FtsK/SpoIIIE family of DNA translocation motors in which the N-terminus forms a hexameric assembly in the IM and the C-terminal cytoplasmic domain has a conserved Walker A motif^43–45^. The hexameric C-terminal domain interacts with the relaxosome complex. DotB is potentially a counterpart of the VirB11 ATPase of the T4ASS. Purified DotB forms hexameric rings in an ATP-independent manner^46^. DotB is more closely related to PilT, an ATPAse of the type IV pilus system, than to VirB11, suggesting a possible origin from the T2SS^47^ Based on previous results^19,42^, we hypothesized that the barrel structures we saw in the T4BSS were the ATPases. Consistent with our hypothesis, all cytoplasmic densities were missing in the ΔdotLOB mutant (Fig. 3S, Y). Given that the size of the tall peripheral barrels matches the known structure of DotO, but is substantially taller than DotL, and DotB has the shortest sequence of all three, we propose that DotO forms the peripheral barrels.

#### DotI/DotJ

DotI and DotJ are closely-related IM proteins that together form a heterocomplex that associates with the Dot/Icm core complex^17^, with DotI essential for Dot/Icm function^11,17,48^. The C-terminal domain of DotI shows structural similarity to VirB8^17^. DotI, VirB8, and TraM (the DotI ortholog in the conjugative R64 plasmid system) all form oligomeric rings in crystals with biologically relevant interfaces, indicating that they likely form rings *in vivo*^17,49^. Although the exact symmetry of the DotIJ ring is not known, Low et al. estimated 12 copies of VirB8 per VirB_3-10_ complex. Based on this, it was suggested that VirB8 might form an IM-associated ring in the periplasm above the hexameric ATPases^17,19^. In the *ΔdotIJ* mutant (Fig. 3T, Z), several differences were apparent: i) the cytoplasmic barrels were missing, ii) the IM was more bent (adopting a shallow V-shape pointing up to the stalk (Fig. S11A-H), iii) the distance between the inner and outer membranes increased from ~31 to 34 nm (center-to-center), and iv) a faint density at the base of the wings was missing. The shape differences in the membrane and vertical expansion of the complex introduced both positive and negative differences on either side of the membrane and between the O- and I-layers, but these cannot correspond to the periplasmic domain of DotI because the linker between its transmembrane and folded domains is not long enough (27 residues: ~9.5 nm max). Instead, we hypothesized that the faint density at the base of the rings was DotI. To consider this more carefully we inspected sub-tomogram averages of wild-type, DotF-sfGFP, and *ΔdotIJ* aligned on the IM (Fig. S11I-N). In both the wild-type and DotF-sfGFP averages there was a ring of density just above the IM at a radius of 8.3 nm from the channel axis, but this was missing in the *ΔdotIJ* map (Fig. S11K, N).

A structure of an octameric complex of DotI periplasmic domains has been solved, as well as several structures of VirB8^17,50,51^. While the curvature of the DotI complex as crystallized did not match the ring curvature we saw *in vivo*, the monomer could be reasonably accommodated within the ring density (Fig. 4), and we suspect that the presence of DotJ likely influences the curvature of the ring. We propose that the periplasmic domain of DotJ forms a 16.6 nm-wide ring around the stalk channel ~3.5 nm above the IM (Fig. S6D), where it stabilizes the ATPases (which are lost in DotIJ’s absence). This is consistent with previous reports that DotIJ forms a ring around the substrate translocation channel^49^.

## Discussion

Here we visualized the structure of the Dot/Icm T4BSS at “macromolecular” (~2-4 nm) resolution and dissected it, pinpointing the location of all 10 proteins with periplasmic domains. We found that the alpha, beta, and gamma rings are 13-fold symmetric. This is surprising because all reported T4ASSs, including those of the R388 and pK101 plasmids and the *H. pylori* Cag T4SS, exhibit 14-fold symmetry^18–21,52^. We also discovered a wide central channel directly above a central cytoplasmic ATPase. A model of the VirB(_3−10_) T4ASS complex from the R388 plasmid showed an OM-associated core complex connected to a cytoplasmic complex by an apparently solid stalk^19^. Similarly, in a reconstruction of the purified *H. pylori* Cag core complex^18^, and in our own *in situ* structure of the complete Cag T4SS^21^, the stalk appears less dense than other features of the particle, as if disordered, but there is no indication of a central lumen. Based on these observations, it was concluded that the T4SS is unlike other secretion systems in that it lacks a central channel^14,19,20,53,54^, leaving open the question of how substrates traverse the machine. Here we saw that the Dot/Icm stalk is a wide channel that could potentially passage substrates. We suggest that the walls of this channel are β-helices formed by a non-conserved sequence of repeats in DotG. It remains unclear whether T4ASSs also have stalk channels or if T4BSSs elaborate pili like T4ASSs. One possibility is that the DotG β-helices play the same role in T4BSSs that pili play in T4ASSs.

Other architectural elements seem to be conserved across the T4SS family. The C-termini of DotG/VirB10, DotH/VirB9, and DotD/VirB7 apparently form an OM complex (O-layer) present in both T4ASSs and T4BSSs. DotC and DotH form another ring just below the OM complex that is likely the counterpart of the I-layer. Finally, we observed that DotI forms a ring around the channel near the IM, just as has been postulated for VirB8^49^. Together DotG, DotH, DotD, and DotC enclose a secretion chamber at the top of the stalk channel. Substrates are likely first secreted through the inner membrane by the short central ATPase positioned directly underneath the channel, then pass through the channel to the secretion chamber, and ultimately exit through a pore in the outer membrane.

There may be an additional pathway for substrates into the machine, however, through the gap between the beta and gamma rings. While there is some density connecting those rings (the DotH elbow), it is much fainter than the rings, suggesting it is not circumferentially continuous (there are holes). In type II and type III secretion systems, the C-terminal domain of PilP/GspC interacts with effectors and with the N0 domain of secretin^55,56^. These effector interactions are high specificity but low affinity, allowing PilP/GspC to recruit effectors from the periplasm and deliver them to the translocation channel. Here, we located DotD, which contains an N0 secretin domain, at the periphery of the O-layer, near the gap between the beta and gamma densities. We found the C-terminal domain of DotF, which is similar to the C-terminal domain of PilP/GspC, at the elbow, just below DotD. In addition, the wing density we identified as DotF appears directly across the IM from the peripheral ATPase barrels. Together this suggests the intriguing possibility that DotF may play a role in translocating effectors secreted into the periplasm into the secretion chamber through the gap between beta and gamma. Alternatively, DotF may play a role in stabilizing the apparatus or triggering conformational changes.

This DotF-elbow interaction would rationalize the observation that DotF recruitment to the poles depends on the presence of DotCDH (the beta and gamma rings)^23^. It may also explain the increased stability of the complex we observed in the DotF-sfGFP mutant: the fusion might stabilize DotF’s interaction with DotD in the elbow, in turn rigidifying everything else from the DotK linkage to the cell wall to the stalk channel walls and even the arrangement of the cytoplasmic ATPases (Fig. S1, S2).

Together with recent immunofluorescence data^23^, our results suggest that assembly occurs in a central-to-peripheral pattern, starting with a seed DotU/IcmF polarization factor that sits in the center of the complex on its axis, followed by assembly of a DotCDH complex around this seed, followed by other components above, below, and to the sides. This is not surprising, since one can imagine evolution adding components to the surface of an existing machine. A primordial DotG/VirB10-like protein alone may have formed a complete channel, spanning all the way from the inner to the outer membrane. Later it might have been stabilized by accretion of a DotH/VirB9-like protein around it, followed by a DotD/VirB7-like protein. An ATPase may have increased the efficiency and specificity of secretion into the DotG/VirB10 channel (and provided an energy source for powered secretion). The capacity to secrete additional effectors may have come with the accretion of additional ATPases on the side (DotO/VirB4), and a periplasmic shuttle system (DotF) to get them into the DotG channel. Supporting this scenario, we found that absence of DotL, DotO, and DotB does not affect assembly of the OM-associated or periplasmic complexes (Fig. 3S), suggesting that these complexes assemble independently of the ATPases. An intriguing obeservaton was that in the *ΔdotIJ* mutant, there were no cytoplasmic densities. Previously, it was suggested that VirB8 might form an IM-associated ring in the periplasm above the hexameric ATPases^17,19^. Our results support that claim and reveal that the DotIJ/VirB8 ring plays a critical role in recruiting the cytoplasmic ATPases.

While many components have homologs in other systems, DotC seems unique to the T4BSSs, which need to be polar for effective Dot/Icm function^57^. DotC may therefore have been acquired on the surface of DotH to mediate targeting to a putatively already-existing polarization factor. DotK could have accreted to further stabilize the system and anchor it to the cell wall.

Jeong et al. recently showed that DotC and DotH form an initial complex, but DotH only associates with the OM after the arrival of DotD^23^. Related to that finding, here we failed to find T4SS-like particles in the *ΔdotD* mutant. Together this might be explained if one or a few DotCH complexes were recruited to each of potentially many individual DotU/IcmF seeds at the pole, but complete rings with 13 copies of DotCH didn’t form until DotD arrives to stabilize the rings by binding around the periphery. Before DotD arrives, individual DotCH complexes would have only one lipoprotein tether to the OM per complex. Complete DotCDH rings would increase the number of lipoprotein links to 26 (13 each from DotCs and DotDs). DotK binding to the periphery of the DotCDH ring would add another 13 lipoprotein links, potentially explaining DotH’s strong OM association. Given its N-terminal association with the IM, DotG may assemble simultaneously with the DotCH ring.

All these ideas and the many claims and relationships presented by the architectural map should open the door to a new era of specific hypothesis testing in the T4SS field and guide efforts towards higher-resolution structure determination of this fascinating and complex cellular nanomachine.

## Materials and Methods

### Plasmid and strain construction

Bacterial strains, plasmids, and primers employed in this study are summarized in SI Table 2. pJB4417, the *dotD dotC dotH* expressing plasmid, was constructed by amplifying *dotD dotC* using primers JVP1591/JVP1594. The PCR product was digested with KpnI and cloned into KpnI-digested pJB1555 *(dotH* complementing clone). pJB1555 was constructed by amplifying *dotH* using primers JPV575/JVP576. The PCR product was digested with KpnI/BamHI and cloned into KpnI-BamHI-digested pJB908. pJB5184, the *ΔdotH ΔdotG ΔdotF*, suicide plasmid, was constructed by amplifying DNA flanking the locus using primers. The PCR fragments were digested with SalI/BamHI and BamHI/NotI, respectively, and cloned into SalI/NotI digested pSR47S. pJB5185, the *ΔdotE ΔdotP* suicide plasmid, was constructed by amplifying DNA flanking the locus using primers JVP375/JVP376 and JVP636/JVP382. The PCR fragments were digested with SalI/BamHI and BamHI/NotI, respectively, and cloned into SalI/NotI digested pSR47S. pJB6162, the *ΔdotJ ΔdotI* suicide plasmid, was constructed by amplifying DNA flanking the locus using primers JVP447/JVP2378 and JVP893/JVP455. The PCR fragments were digested with SalI/BamHI and BamHI/NotI, respectively, and cloned into SalI/NotI digested pSR47S.

JV6781, the triple ATPase (*dotL, dotO, dotB*) mutant, was constructed by deleting *dotO* in JV5629 (*ΔdotL ΔdotB*) using the *dotO* suicide plasmid pJB1333. JV5629 was constructed by deleting *dotL* in JV918 *(ΔdotB)* using the *dotL* suicide plasmid pJB1001. pJB7255, the DotF-sfGFP integration plasmid, was constructed by amplifying DotF-sfGFP (using JVP2973/JVP2990) and *dotE* (using JVP2992/JVP2993). The PCR fragments were digested with BamHI/NotI and NotI/SacI, respectively, and cloned into BamHI/SacI digested pSR47S. JV7058, the *ΔdotH ΔdotG ΔdotF* strain, was constructed by integration of pJB5184 into Lp02 followed by resolution of the merodiploid. JV7091, the *ΔdotE ΔdotP* strain, was constructed by integration of pJB5185 into Lp02 followed by resolution of the merodiploid. JV7967, the *ΔdotJ ΔdotI* strain, was constructed by integration of pJB6162 into Lp02 followed by resolution of the merodiploid. JV9082, the DotF-sfGFP chromosomally integrated strain, was constructed by integration of pJB7255 into Lp02 followed by resolution of the merodiploid. JV9114, the DotC-sfGFP chromosomally integrated strain, was constructed in two steps. First, a SacB/CmR cassette was integrated prior to the stop codon of DotC using pJB7264 by natural transformation of Lp02. Then the cassette was replaced in a second natural transformation step using pJB7283.

### Western blot analysis

Protein samples were boiled for 5 min in Laemmli sample buffer and separated by sodium dodecyl sulfate–polyacrylamide gel electrophoresis, followed by transfer to polyvinylidene difluoride membranes. Membranes were blocked in BLOTTO (PBS containing 5% non-fat dry milk), washed with wash buffer (PBS containing 0.05% Tween 20) and incubated for 1 hour with antibody diluted in BLOTTO. Blots were then washed with wash buffer followed by 1 hour incubation with secondary goat anti-rabbit antibody conjugated to horseradish peroxidase (Sigma) diluted 1:10,000 in BLOTTO. Blots were subsequently washed with wash buffer prior to development using an ECL detection kit (GE Healthcare).

### Sample preparation for electron cryotomography

*L. pneumophila* Lp02 cells were harvested at early stationary phase (OD600 of ~3.0), mixed with 10-nm colloidal gold beads (Sigma-Aldrich, St. Louis, MO) precoated with bovine serum albumin, and applied onto freshly glow-discharged copper R2/2 200 Quantifoil holey carbon grids (Quantifoil Micro Tools GmbH, Jena, Germany). Grids were then blotted and plunge-frozen in a liquid ethane/propane mixture^58^ using an FEI Vitrobot Mark IV and stored in liquid nitrogen for subsequent imaging.

### Electron cryotomography, sub-tomogram averaging, and difference analysis

Tilt-series were recorded of frozen *L. pneumophila* Lp02 cells in an FEI Titan Krios 300 kV field emission gun transmission electron microscope (FEI Company, Hillsboro, OR) equipped with a Gatan imaging filter (Gatan, Pleasanton, CA) and a K2 Summit direct detector in counting mode (Gatan, Pleasanton, CA) using the UCSF Tomography software^59^ and a total dose of ~ 100 e/A^2^ per tilt-series and target defocus of ~6 μm underfocus. Images were aligned, contrast transfer function corrected, and reconstructed using IMOD^60^. SIRT reconstructions were produced using TOMO3D^61^ and sub-tomogram averaging was performed using PEET^62^. Finally, the local resolution was calculated by ResMap^63^. As the Dot/Icm sub-tomogram average exhibited a gross two-fold symmetry around the central mid-line in the periplasm, we applied two-fold symmetry in those regions to produce the 2-D figures shown, but no symmetry was applied to the cytoplasmic densities due to their poor resolution. Sub-tomograms were aligned within masks centered either on the OM-associated densities, the IM-associated densities, or the densities between the IM and the gamma ring. The figures are composites, showing the average that was clearest in that region of the T4BSS, with lines visible at the interface to remind the viewer of this fact. Difference maps were created by mutually aligning two averages, then subtracting the densities of the mutant from the densities of the reference (usually the wildtype). Losses and increases of density were shown in yellow and red, respectively, and shaded according to significance (bright and light color for differences greater than two and one standard deviations above the mean, respectively).

### Structure prediction and model building

Gene sequences were obtained from UniProt^64^. Signal sequences were predicted with SignalP 4.1^65^. Transmembrane domains were predicted by Phobius^66^ and TMHMM Server^67^. Domain structures were predicted by servers Phyre2^68^, I-TASSER^69^, and Quark^70^. The cysteines attached to the OM in lipoproteins were marked as reported in the literature or predicted using SignalP. Candidate ring structures with different numbers of monomers were generated using SymmDock^71^ (for instance the ring of DotG N-terminal β-helices).

## Acknowledgments

This work was supported by NIH grant R01AI127401 to G.J.J. Some data were recorded at the cryo-EM facility at Janelia Research Campus. We thank Dr. Catherine Oikonomou for creation of the domain maps (Figs. S4, S5) and for help structuring and revising the text. We thank Jacob Gyore for running the Western blots in Fig. S3.

## References

1. Costa, T. R. D. et al. Secretion systems in Gram-negative bacteria: structural and mechanistic insights. Nat. Rev. Microbiol. 13, 343–359 (2015).

2. Green, E. R. & Mecsas, J. Bacterial Secretion Systems: An Overview. Microbiol. Spectr. 4, (2016).

3. Censini, S., Stein, M. & Covacci, A. Cellular responses induced after contact with Helicobacter pylori. Curr. Opin. Microbiol. 4, 41–46 (2001).

4. Backert, S. & Meyer, T. F. Type IV secretion systems and their effectors in bacterial pathogenesis. Curr. Opin. Microbiol. 9, 207–217 (2006).

5. Christie, P. J. & Vogel, J. P. Bacterial type IV secretion: conjugation systems adapted to deliver effector molecules to host cells. Trends Microbiol. 8, 354–360 (2000).

6. Guglielmini, J. et al. Key components of the eight classes of type IV secretion systems involved in bacterial conjugation or protein secretion. Nucleic Acids Res. 42, 5715–5727 (2014).

7. Christie, P. J., Atmakuri, K., Krishnamoorthy, V., Jakubowski, S. & Cascales, E. Biogenesis, architecture, and function of bacterial type IV secretion systems. Annu. Rev. Microbiol. 59, 451–485 (2005).

8. Christie, P. J., Whitaker, N. & González-Rivera, C. Mechanism and structure of the bacterial type IV secretion systems. Biochim. Biophys. Acta 1843, 1578–1591 (2014).

9. Chandran Darbari, V. & Waksman, G. Structural Biology of Bacterial Type IV Secretion Systems. Annu. Rev. Biochem. 84, 603–629 (2015).

10. Segal, G., Feldman, M. & Zusman, T. The Icm/Dot type-IV secretion systems of Legionella pneumophila and Coxiella burnetii. FEMS Microbiol. Rev. 29, 65–81 (2005).

11. Segal, G., Purcell, M. & Shuman, H. A. Host cell killing and bacterial conjugation require overlapping sets of genes within a 22-kb region of the Legionella pneumophila genome. Proc. Natl. Acad. Sci. U. S. A. 95, 1669–1674 (1998).

12. Vogel, J. P., Andrews, H. L., Wong, S. K. & Isberg, R. R. Conjugative transfer by the virulence system of Legionella pneumophila. Science 279, 873–876 (1998).

13. Nagai, H. & Kubori, T. Type IVB Secretion Systems of Legionella and Other Gram-Negative Bacteria. Front. Microbiol. 2, 136 (2011).

14. Ghosal, D., Chang, Y.-W., Jeong, K. C., Vogel, J. P. & Jensen, G. J. In situ structure of the Legionella Dot/Icm type IV secretion system by electron cryotomography. EMBO Rep. 18, 726–732 (2017).

15. Nakano, N., Kubori, T., Kinoshita, M., Imada, K. & Nagai, H. Crystal structure of Legionella DotD: insights into the relationship between type IVB and type II/III secretion systems. PLoS Pathog. 6, e1001129 (2010).

16. Kwak, M.-J. et al. Architecture of the type IV coupling protein complex of Legionella pneumophila. Nat. Microbiol. 2, 17114 (2017).

17. Kuroda, T. et al. Molecular and structural analysis of Legionella DotI gives insights into an inner membrane complex essential for type IV secretion. Sci. Rep. 5, (2015).

18. Frick-Cheng, A. E. et al. Molecular and Structural Analysis of the *Helicobacter pylori cag* Type IV Secretion System Core Complex. mBio 7, e02001–15 (2016).

19. Low, H. H. et al. Structure of a type IV secretion system. Nature 508, 550–553 (2014).

20. Chandran, V. et al. Structure of the outer membrane complex of a type IV secretion system. Nature 462, 1011–1015 (2009).

21. Chang, Y.-W., Shaffer, C. L., Rettberg, L. A., Ghosal, D. & Jensen, G. J. In Vivo Structures of the Helicobacter pylori cag Type IV Secretion System. Cell Rep. 23, 673–681 (2018).

22. Vincent, C. D. et al. Identification of the core transmembrane complex of the Legionella Dot/Icm type IV secretion system. Mol. Microbiol. 62, 1278–1291 (2006).

23. Jeong, K. C., Gyore, J., Teng, L., Ghosal, D., Jensen, G. J. & Vogel, J. P. Polar targeting and assembly of the Legionella Dot/Icm type IV secretion system (T4SS) by co-option of T6SS components (manuscript in preparation).

24. Morozova, I. Comparative sequence analysis of the icm/dot genes in Legionella. Plasmid 51, 127–147 (2004).

25. Segal, G. & Shuman, H. A. Legionella pneumophila utilizes the same genes to multiply within Acanthamoeba castellanii and human macrophages. Infect. Immun. 67, 2117–2124 (1999).

26. Rivera-Calzada, A. et al. Structure of a bacterial type IV secretion core complex at subnanometre resolution. EMBO J. 32, 1195–1204 (2013).

27. Souza, D. P. et al. A component of the Xanthomonadaceae type IV secretion system combines a VirB7 motif with a N0 domain found in outer membrane transport proteins. PLoS Pathog. 7, e1002031 (2011).

28. Sexton, J. A., Miller, J. L., Yoneda, A., Kehl-Fie, T. E. & Vogel, J. P. Legionella pneumophila DotU and IcmF Are Required for Stability of the Dot/Icm Complex. Infect. Immun. 72, 5983–5992 (2004).

29. Cascales, E. The type VI secretion toolkit. EMBO Rep. 9, 735–741 (2008).

30. Durand, E. et al. Structural characterization and oligomerization of the TssL protein, a component shared by bacterial type VI and type IVb secretion systems. J. Biol. Chem. 287, 14157–14168 (2012).

31. Durand, E. et al. Biogenesis and structure of a type VI secretion membrane core complex. Nature 523, 555–560 (2015).

32. Matthews, M. & Roy, C. R. Identification and subcellular localization of the Legionella pneumophila IcmX protein: a factor essential for establishment of a replicative organelle in eukaryotic host cells. Infect. Immun. 68, 3971–3982 (2000).

33. Nagai, H. & Roy, C. R. The DotA protein from Legionella pneumophila is secreted by a novel process that requires the Dot/Icm transporter. EMBO J. 20, 5962–5970 (2001).

34. Luedtke, B. E., Mahapatra, S., Lutter, E. I. & Shaw, E. I. The Coxiella Burnetii type IVB secretion system (T4BSS) component DotA is released/secreted during infection of host cells and during in vitro growth in a T4BSS-dependent manner. Pathog. Dis. 75, (2017).

35. Maneewannakul, S., Maneewannakul, K. & Ippen-Ihler, K. Characterization, localization, and sequence of F transfer region products: the pilus assembly gene product TraW and a new product, TrbI. J. Bacteriol. 174, 5567–5574 (1992).

36. Sutherland, M. C., Binder, K. A., Cualing, P. Y. & Vogel, J. P. Reassessing the role of DotF in the Legionella pneumophila type IV secretion system. PloS One 8, e65529 (2013).

37. Luo, Z.-Q. & Isberg, R. R. Multiple substrates of the Legionella pneumophila Dot/Icm system identified by interbacterial protein transfer. Proc. Natl. Acad. Sci. U. S. A. 101, 841–846 (2004).

38. Christie, P. J. The Mosaic Type IV Secretion Systems. EcoSal Plus 7, (2016).

39. Kirby, J. E., Vogel, J. P., Andrews, H. L. & Isberg, R. R. Evidence for pore-forming ability by Legionella pneumophila. Mol. Microbiol. 27, 323–336 (1998).

40. Coers, J. et al. Identification of Icm protein complexes that play distinct roles in the biogenesis of an organelle permissive for Legionella pneumophila intracellular growth. Mol. Microbiol. 38, 719–736 (2000).

41. Roy, C. R. & Isberg, R. R. Topology of Legionella pneumophila DotA: an inner membrane protein required for replication in macrophages. Infect. Immun. 65, 571–578 (1997).

42. Peña, A. et al. The hexameric structure of a conjugative VirB4 protein ATPase provides new insights for a functional and phylogenetic relationship with DNA translocases. J. Biol. Chem. 287, 39925–39932 (2012).

43. Aussel, L. et al. FtsK Is a DNA motor protein that activates chromosome dimer resolution by switching the catalytic state of the XerC and XerD recombinases. Cell 108, 195–205 (2002).

44. Errington, J., Bath, J. & Wu, L. J. DNA transport in bacteria. Nat. Rev. Mol. Cell Biol. 2, 538–545 (2001).

45. Massey, T. H., Mercogliano, C. P., Yates, J., Sherratt, D. J. & Löwe, J. Double-stranded DNA translocation: structure and mechanism of hexameric FtsK. Mol. Cell 23, 457–469 (2006).

46. Savvides, S. N. et al. VirB11 ATPases are dynamic hexameric assemblies: new insights into bacterial type IV secretion. EMBO J. 22, 1969–1980 (2003).

47. Sexton, J. A. et al. The Legionella pneumophila PilT homologue DotB exhibits ATPase activity that is critical for intracellular growth. J. Bacteriol. 186, 1658–1666 (2004).

48. Andrews, H. L., Vogel, J. P. & Isberg, R. R. Identification of linked Legionella pneumophila genes essential for intracellular growth and evasion of the endocytic pathway. Infect. Immun. 66, 950–958 (1998).

49. Gillespie, J. J. et al. Structural Insight into How Bacteria Prevent Interference between Multiple Divergent Type IV Secretion Systems. mBio 6, e01867–01815 (2015).

50. Bailey, S., Ward, D., Middleton, R., Grossmann, J. G. & Zambryski, P. C. Agrobacterium tumefaciens VirB8 structure reveals potential protein-protein interaction sites. Proc. Natl. Acad. Sci. U. S. A. 103, 2582–2587 (2006).

51. Terradot, L. et al. Structures of two core subunits of the bacterial type IV secretion system, VirB8 from Brucella suis and ComB10 from Helicobacter pylori. Proc. Natl. Acad. Sci. U. S. A. 102, 4596–4601 (2005).

52. Fronzes, R. et al. Structure of a Type IV Secretion System Core Complex. Science 323, 266–268 (2009).

53. Hu, B., Lara-Tejero, M., Kong, Q., Galán, J. E. & Liu, J. In Situ Molecular Architecture of the Salmonella Type III Secretion Machine. Cell 168, 1065–1074.e10 (2017).

54. Chang, Y., Rettberg, L. A., Ortega, D. R. & Jensen, G. J. *In vivo* structures of an intact type VI secretion system revealed by electron cryotomography. EMBO Rep. 18, 1090–1099 (2017).

55. Douzi, B. et al. Unraveling the Self-Assembly of the Pseudomonas aeruginosa XcpQ Secretin Periplasmic Domain Provides New Molecular Insights into Type II Secretion System Secreton Architecture and Dynamics. mBio 8, (2017).

56. Nivaskumar, M. & Francetic, O. Type II secretion system: A magic beanstalk or a protein escalator. Biochim. Biophys. Acta BBA - Mol. Cell Res. 1843, 1568–1577 (2014).

57. Yerushalmi, G., Zusman, T. & Segal, G. Additive effect on intracellular growth by Legionella pneumophila Icm/Dot proteins containing a lipobox motif. Infect. Immun. 73, 7578–7587 (2005).

58. Tivol, W. F., Briegel, A. & Jensen, G. J. An improved cryogen for plunge freezing. Microsc. Microanal. Off. J. Microsc. Soc. Am. Microbeam Anal. Soc. Microsc. Soc. Can. 14, 375–379 (2008).

59. Zheng, S. Q. et al. UCSF tomography: an integrated software suite for real-time electron microscopic tomographic data collection, alignment, and reconstruction. J. Struct. Biol. 157, 138–147 (2007).

60. Kremer, J. R., Mastronarde, D. N. & McIntosh, J. R. Computer visualization of three-dimensional image data using IMOD. J. Struct. Biol. 116, 71–76 (1996).

61. Agulleiro, J.-I. & Fernandez, J.-J. Tomo3D 2.0--exploitation of advanced vector extensions (AVX) for 3D reconstruction. J. Struct. Biol. 189, 147–152 (2015).

62. Nicastro, D. et al. The molecular architecture of axonemes revealed by cryoelectron tomography. Science 313, 944–948 (2006).

63. Kucukelbir, A., Sigworth, F. J. & Tagare, H. D. Quantifying the local resolution of cryo-EM density maps. Nat. Methods 11, 63–65 (2014).

64. UniProt Consortium, T. UniProt: the universal protein knowledgebase. Nucleic Acids Res. 46, 2699 (2018).

65. Emanuelsson, O., Brunak, S., von Heijne, G. & Nielsen, H. Locating proteins in the cell using TargetP, SignalP and related tools. Nat. Protoc. 2, 953–971 (2007).

66. Käll, L., Krogh, A. & Sonnhammer, E. L. L. A combined transmembrane topology and signal peptide prediction method. J. Mol. Biol. 338, 1027–1036 (2004).

67. Krogh, A., Larsson, B., von Heijne, G. & Sonnhammer, E. L. Predicting transmembrane protein topology with a hidden Markov model: application to complete genomes. J. Mol. Biol. 305, 567–580 (2001).

68. Kelley, L. A., Mezulis, S., Yates, C. M., Wass, M. N. & Sternberg, M. J. E. The Phyre2 web portal for protein modeling, prediction and analysis. Nat. Protoc. 10, 845–858 (2015).

69. Yang, J. et al. The I-TASSER Suite: protein structure and function prediction. Nat. Methods 12, 7–8 (2015).

70. Xu, D. & Zhang, Y. Ab initio protein structure assembly using continuous structure fragments and optimized knowledge-based force field. Proteins 80, 1715–1735 (2012).

71. Schneidman-Duhovny, D., Inbar, Y., Nussinov, R. & Wolfson, H. J. PatchDock and SymmDock: servers for rigid and symmetric docking. Nucleic Acids Res. 33, W363–367 (2005).

72. Jeong, K. C., Ghosal, D., Chang, Y.-W., Jensen, G. J. & Vogel, J. P. Polar delivery of Legionella type IV secretion system substrates is essential for virulence. Proc. Natl. Acad. Sci. U. S.A. 114, 8077–8082 (2017).

